# Expression Strategies for Recombinant HECT E3 Ligases in *Escherichia coli*

**DOI:** 10.1101/2023.12.28.573528

**Authors:** Paulina Krach, Daria Gajewska, Maria Sagan, Anna Szlachcic, Michał J. Walczak

## Abstract

A comparative analysis of recombinant expression in *E. coli* of three HECT E3 ligases reveals a consistent preference for lower expression temperatures. Lower temperatures prevent aggregation and misfolding, enhancing the efficiency and solubility of expressed HECT ligases. Isolated HECT domains generally exhibit higher expression success compared to full-length counterparts, offering improved solubility and yields. However, expression levels vary among ligases, necessitating tailored strategies. Future studies may explore full-length HECT-type E3 ligases in covalent complexes with ubiquitin as a potential, generalizable platform for biomedical research.

## Introduction

Ubiquitination is a post-translational modification (PTM) in which ubiquitin, a 76-amino acid protein, is covalently attached to target protein (Hershko & Ciechanover, 1998; Grabbe *et al*., 2011). This process plays a crucial role in regulating various cellular functions, and its significance is underscored by its involvement in several key biological processes. Ubiquitination marks proteins for degradation, identifying and eliminating misfolded or damaged proteins, but also regulates cell cycle, modulates signaling pathways, and influences development and differentiation (Sun & Chen, 2004; Komander, 2009). It is also essential for preventing diseases, as the uncontrolled accumulation of misfolded proteins leads to cellular dysfunction (Popovic *et al*., 2014; Atkin & Paulson, 2014).

E3 ligases play a central role in the ubiquitination process, as they are the proteins responsible for transferring ubiquitin molecules from ubiquitin-conjugating enzymes to the target protein. There are three main types of enzymes involved in the ubiquitination process: E1 (ubiquitin-activating enzyme), E2 (ubiquitin-conjugating enzyme), and E3 (ubiquitin ligase) (Hershko & Ciechanover, 1998).

The specificity of ubiquitination is largely determined by the interaction between the E3 ligase and its substrate. Various E3 ligases recognize specific target proteins and facilitate the transfer of ubiquitin to these targets. This specificity ensures that ubiquitin is attached to the right proteins at the right time. This is reflected in the diversity of E3 ligases: there is more than 600 E3 ligases with distinct structural and functional characteristics. This diversity enables a broad spectrum of substrates to be targeted for ubiquitination, thereby contributing to the regulation of various cellular processes (Berndsen & Wolberger, 2014; Zheng & Shabek, 2017).

Another aspect is the recently emerging therapeutic strategies utilizing E3 ligases, such as targeted protein degradation (TPD). TPD approaches leverage E3 ligases to selectively degrade specific proteins of interest (Dong *et al*., 2021; Cowan & Ciulli, 2022). This precision targeting enables the development of therapies that can specifically eliminate disease-causing or aberrant proteins while sparing normal cellular proteins. The completely different mode of action enables the targeting of proteins that were previously considered undruggable by traditional small-molecule inhibitors (Cromm & Crews, 2017; Bekes *et al*., 2022).

The E3 ligases most commonly employed in such therapeutic approaches belong to the RING (Really Interesting New Gene) family of E3 ligases, with cereblon (CRBN) and von Hippel-Lindau (VHL) ligases thoroughly characterized. However, the field is expanding its toolbox and exploring the possibility of additional E3 ligases for enhanced therapeutic applications (Schapira *et al*., 2019).

HECT (Homologous to E6-AP Carboxyl Terminus) E3 ligases contain a catalytic C-terminal HECT domain, and the substrate specificity of the ligase is determined by its variable N-terminal part (Rotin & Kumar, 2009). HECT E3 ligases are much less studied than RING ligases, partially due to the more complex structure compared to RING ligases coupled with scarcity of high-resolution structural information for many of them. This hinders detailed insights into their mechanisms and interactions, making it harder to study them experimentally. Engagement of HECT E3 ligases into TPD technology has also not been demonstrated, potentially due to differences in their mechanism of action (MoA) compared to RING ones. Unlike RING ligases, which catalyze transfer of ubiquitin from E2 conjugating enzyme to a substrate, HECT ligases first undergo ubiquitination and dissociation from E2. Next, they recognize substrates to which they attach ubiquitin chains, which makes them a *bona fide* catalysts in contrast to RING E3s. This unique mechanism makes HECT ligases attractive proteins for the degradation technology, especially from the perspective that they do not need multiprotein complex assembly to fulfil their role, and consequently they may form ternary complexes (complexes of E3 ligase:degrader drug:substrate) of favorable geometry and absent of steric hindrances.

In this study, we present an expression screen aimed at identifying optimal conditions for the expression of representative HECT ligase proteins in *Escherichia coli*, with the specific goal of obtaining soluble proteins. The focus is on overcoming challenges associated with the expression and solubility of HECT ligases, essential components of the ubiquitination process. By systematically screening different conditions, we aim to establish a robust protocol for obtaining soluble HECT ligases, facilitating their subsequent characterization.

### Experimental Procedures

#### Plasmid construction

To prepare GST-fused recombinant HECT ligases, codon-optimized sequences of full-length or truncated proteins (Table 1) were introduced into pGEX-6P vector. Constructs were prepared using Gibson assembly strategy, with Gblocks ordered from IDT (Integrated DNA Technology, USA). Competent *E. coli* DH5α cells were transformed with Gibson assembly reaction mix, colonies from each transformation plate were chosen to identify the positive transformants by colony PCR and correct vector and insert sequence was confirmed by DNA sequencing.

#### Protein expression tests

The *E. coli* Bl21 (DE3) strain was transformed with pGEX-6P vectors encoding the full-length or truncated versions of tested E3 ligases: Smurf1 (full-length and HECT domain: aa 420-757), Smurf2 (full-length and HECT domain: aa 414-748) and Itch (full-length). Single colonies were used to inoculate an overnight culture (10mL LB medium supplemented with 50 µg/mL kanamycin) grown at 37C. The overnight culture was used to inoculate (at 1:100 ratio) a 100 mL culture (LB with kanamycin) and the cells were grown until they reached OD_600_ = 0.8. Expression was induced with 0.5 mM Isopropyl β-D-1-thiogalactopyranoside (IPTG), then bacteria were split and cultivated in 3 flasks 25 ml of LB each and cultured at 18C, 25C and 37C. Samples were collected at 2h, 4h and 18h (o/n) post induction. OD_600_ was measured before harvesting the cells. Cells were resuspended in 25 mM phosphate buffer pH 7.8, 200 mM NaCl, 1 mM DTT, 1 mM PMSF and lysed by 3 cycles of sonication. The soluble and insoluble lysate fractions were separated by centrifugation (20 000 g, 10 min), mixed with Laemmli sample loading buffer, heated and loaded on SDS-PAGE normalizing for culture final OD_600_.

#### Small-scale protein purifications

Bacterial pellets corresponding to expression screen conditions with largest amount of soluble protein were lysed in lysis buffer (25 mM phosphate buffer pH 7.8, 200 mM NaCl, 1 mM DTT, 1 mM PMSF) and the soluble fraction separated by centrifugation (20 000 g, 15 min). Supernatant was used for small scale affinity purification – protein of interest was bound to GST-Sepharose beads for 1h at 4C, the resin was washed 3 times with lysis buffer and incubated for 10 min with elution buffer with 10 mM glutathione. Normalized amounts of eluate was analyzed by SDS-PAGE to compare the yields of purified proteins.

## Results and discussion

### Expression of recombinant full-length HECT E3 ligases

Three exemplary E3 ligases from the HECT subfamily**: Smurf1, Smurf2** and **Itch** were cloned as GST-fusions to facilitate expression of recombinant proteins in *E. coli* BL21(DE3) cells. Human HECT ligases can be categorized into three subfamilies—Nedd4, HERC, and other HECTs—based on their N-terminal extensions. The ligases examined in this study belong to the Nedd4 subfamily, characterized by a C2 domain at the N-terminus and a variable number of repeats of tryptophan-tryptophan (WW) motifs (Fig. 1).

**Fig. 1.**
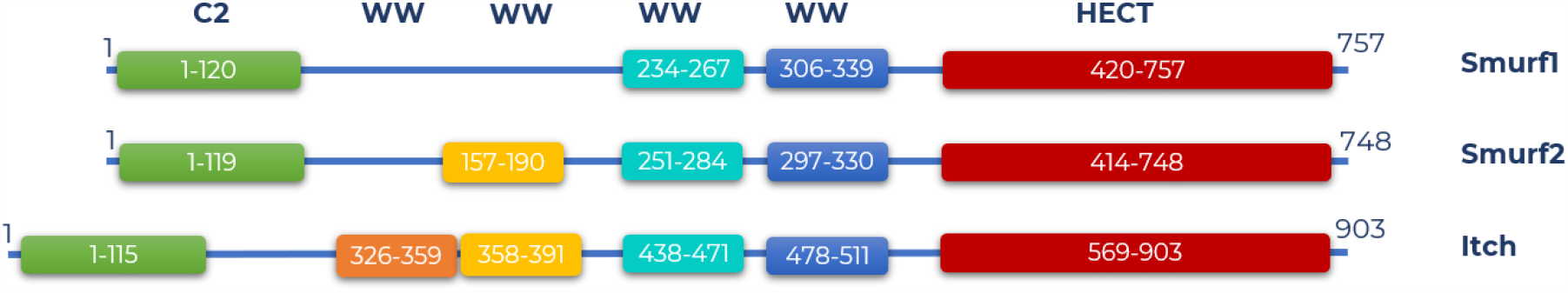
Schematic representation of HECT E3 ligases: Smurf1, Smurf2 and Itch. C2 – C2 domain, WW – Tryptophan-Tryptophan motif..

RING-type E3 ligases are often amenable to expression in *E. coli* as fusion proteins with glutathione S-transferase (GST) tags. Expression of HECT family proteins was reported before as feasible in bacterial systems (Ronchi *et al*., 2014; Hatstata & McCafferty, 2020), therefore the *E. coli*-based system was chosen. In all used constructs codon-optimized coding sequences were used to maximize the chances of obtaining soluble and functional proteins.

Our expression screening approach focused on two main variables – temperature in which the bacteria were cultured, and the post-induction culture time. This classic expression screening setup includes both the optimal for these bacteria temperature of 37***°***C, as well as the low temperature condition (18***°***C), which promotes expression of lower levels of recombinant protein, but with slowed down the rate of protein synthesis minimizing the likelihood of misfolding and aggregation. Low temperature post-induction can benefit from the increased availability of chaperones at lower temperatures, additionally contributing to improved protein solubility (Baneyx *et al*., 2004; Sørensen & Mortensen, 2005; Kaur *et al*., 2018). On the other hand, 37***°***C condition ensures rapid *E. coli* growth, translating to fast biomass production. Temperature of 25***°***C was also included, covering the middle-ground in case the high- and low-temperature points would be too extreme for one of the tested proteins.

Time post-induction is the second parameter that greatly influences the yield of recombinant proteins produced (Kaur *et al*., 2018), however, similarly to optimal expression temperature, it is strongly dependent on the characteristics of the target protein. Therefore, the optimal post-induction time should be determined empirically for each specific case and here the timepoints 2h, 4h and 18h (overnight) were tested.

The initial assessment involved the expression of full-length protein variants (Fig. 2). For GST-Smurf1(FL) majority of the protein is present in the pellet fraction, regardless of the conditions (Fig. 2A). Only overnight culture at 25***°***C showed the presence of recombinant Smurf1(FL) in the soluble portion lysate, and this bacterial pellet was employed for subsequent small-scale purification steps.

**Fig. 2.**
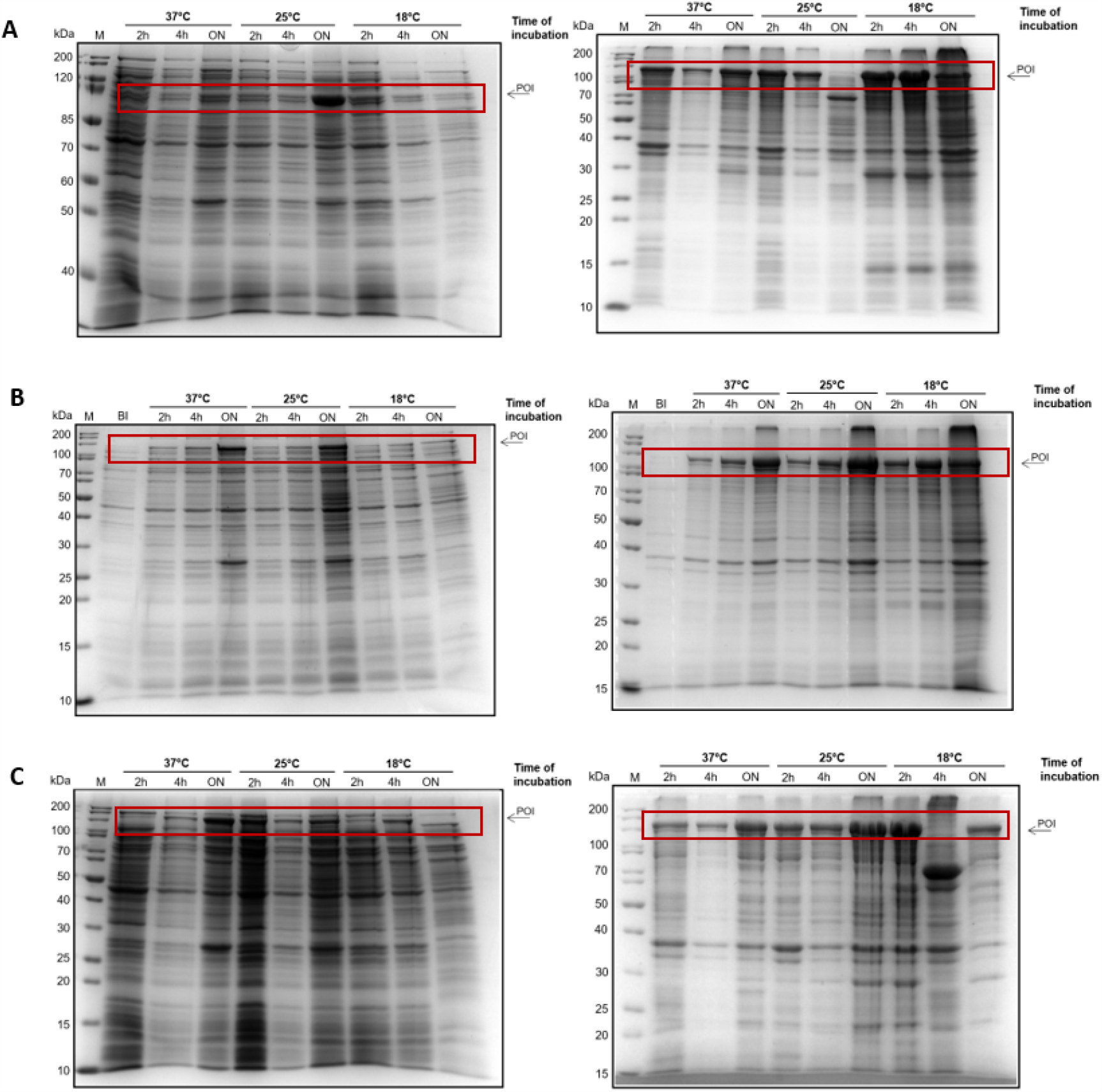
Analysis of GST-Smurf1(FL) (A), GST-Smurf2(FL) (B) and GST-Itch(FL) (C) expression in BL21(DE3) strain. Coomassie Blue staining of 12% polyacrylamide gel after electrophoretic separation of samples, normalized based on the OD_600_ values, of soluble **(left)** and insoluble **(right)** proteins extracted from indicated cultures. Abbreviations: ***M*** – PageRuler Unstained Protein Ladder; ***2h, 4h ON (overnight)*** – time of culture incubation after recombinant protein expression induction with 0.5 mM IPTG; ***37, 25 and 18 C*** – temperature of expression post-induction. Protein molecular weights (kDa) are indicated on the left. Protein of Interest, POI, is indicated with an arrow on the right side of the gel.

For GST-Smurf2(FL), the protein predominantly localized in the insoluble fraction of the bacterial lysate across various tested conditions. However, a discernible trend emerged: the lower the expression temperature and the longer the post-induction time, the greater the proportion of the protein observed in the soluble fraction. Notably, the condition of 18°C with overnight culture exhibited a significant enhancement, with approximately 20% of the expressed protein found in the soluble fraction (Fig. 2B).

Similarly, GST-Itch(FL) demonstrated a tendency towards insolubility, with the majority of the protein residing in the insoluble fraction of the lysate. Only a small fraction of the total protein was detected in the soluble lysate fraction, and the most promising condition proved to be 25°C with long (overnight) expression time (Fig. 2C).

### Small scale-purification of full-length HECT E3 ligases

To quantify the yield of protein achievable under the most promising conditions identified during expression screening and to validate that the bands observed on SDS-PAGE indeed correspond to the Protein of Interest (POI) GST-fusions, we conducted small-scale affinity purifications.

This involved utilizing glutathione resin as part of the purification process, aiming to confirm the specific binding of the GST-fused POI and quantifying the amount of purified protein obtained. This purification step serves to validate the success of the chosen conditions and ensures that the protein visible in the SDS-PAGE analysis of soluble fractions of bacterial lysates can be recovered. It verifies that the protein is not aggregated, and the fusion tag is folded and functional.

As illustrated in Fig. 3, there were notable differences in the purification yields of full-length Smurf1, Smurf2, and Itch ligases. Specifically, purified protein was successfully obtained for Itch(FL), indicating an efficient purification process. However, the yields for Smurf2(FL) were considerably smaller, suggesting potential challenges in the purification of the full-length Smurf2 ligase. Remarkably, no protein eluted for Smurf1(FL), indicating a lack of successful purification under the tested conditions. These differences in purification outcomes highlight distinct behaviors among the three full-length ligases during the purification process.

**Fig. 3.**
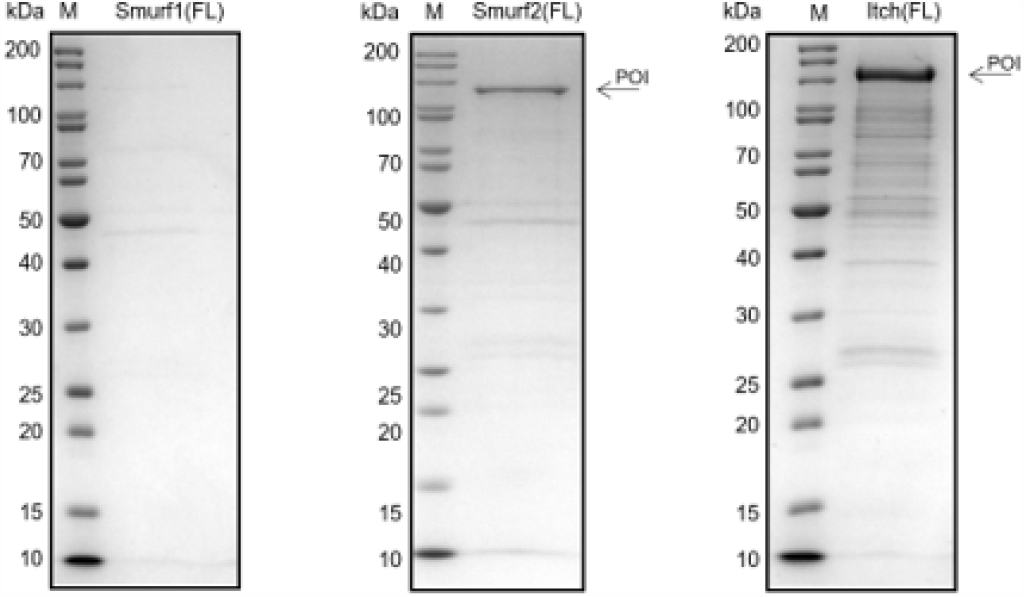
Purification of GST-fusion proteins of Itch(FL) and Smurf2(FL) from bacterial lysates. Soluble fractions of lysates were subjected to affinity-purification and normalized amounts of eluates were analyzed by SDS-PAGE.

Interestingly, the observed efficiency of expression and purification (Itch > Smurf2 > Smurf1) did not correlate with the protein length or the number of WW domains in the N-terminal part of the protein, which distinguishes them from each other (Fig. 1).

### Expression of recombinant isolated HECT domains

Expressing truncated recombinant proteins, as opposed to their full-length counterparts, is a strategy often employed in recombinant protein expression when the expression and/or purification of the full-length version presents challenges [ref]. Truncated proteins may exhibit better solubility compared to their full-length counterparts. Removing certain domains or regions that contribute to aggregation or misfolding can enhance the solubility of the expressed protein, as well as simplify the purification process and therefore increase overall yields.

The HECT domain is the indispensable functional component for the ligases examined in this study. Recognizing its critical role, we opted to investigate the expression of truncated versions of Smurf1 and Smurf2. These truncated versions contain only the HECT domain of the respective ligases. This focused exploration aims to elucidate the expression patterns and behaviors of the HECT domain alone, providing insights into its standalone functionality within each ligase.

The GST-fusion of the Smurf1 HECT domain exhibited limited solubility, similar to the full-length protein. However, under conditions involving lower temperatures and extended expression times (18°C, overnight), the band corresponding to the recombinant protein was detected in the soluble fraction, which underwent subsequent small-scale purification.

Conversely, the HECT domain from Smurf2 demonstrated very high levels of soluble protein expression, especially when bacteria were cultured at 25°C (Fig. 4B). A noticeable trend indicated that with longer expression times, more protein was produced. The protein remained soluble at 18°C and 25°C; however, at the higher temperature of 37°C, longer expression times led to a reduced amount of soluble protein, with a greater presence in the pellet fraction.

**Fig. 4.**
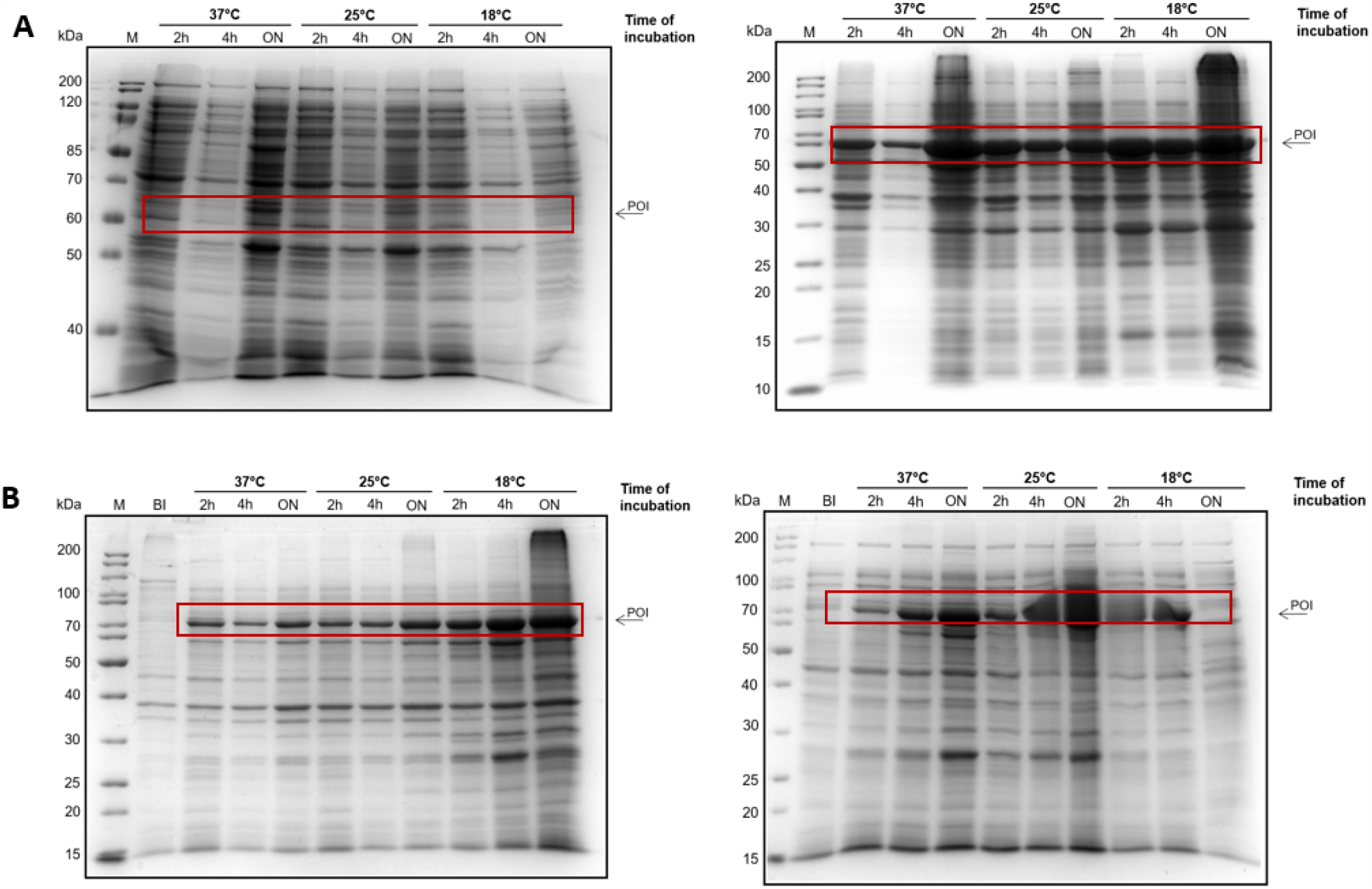
Analysis of GST-Smurf1(420-757) (A) and GST-Smurf2(414-748) expression in BL21(DE3) strain. Coomassie Blue staining of 12% polyacrylamide gel after electrophoretic separation of samples, normalized based on the OD_600_ values, of soluble **(left)** and insoluble **(right)** proteins extracted from indicated cultures. Abbreviations: ***M*** – PageRuler Unstained Protein Ladder; ***2h, 4h ON*** – time of culture incubation after recombinant protein expression induction with 0.5 mM IPTG; ***37°C, 25°C and 18°C*** – temperature of expression post-induction. Protein molecular weights (kDa) are indicated on the left. POI is indicated with an arrow on the right.

**Fig. 5.**
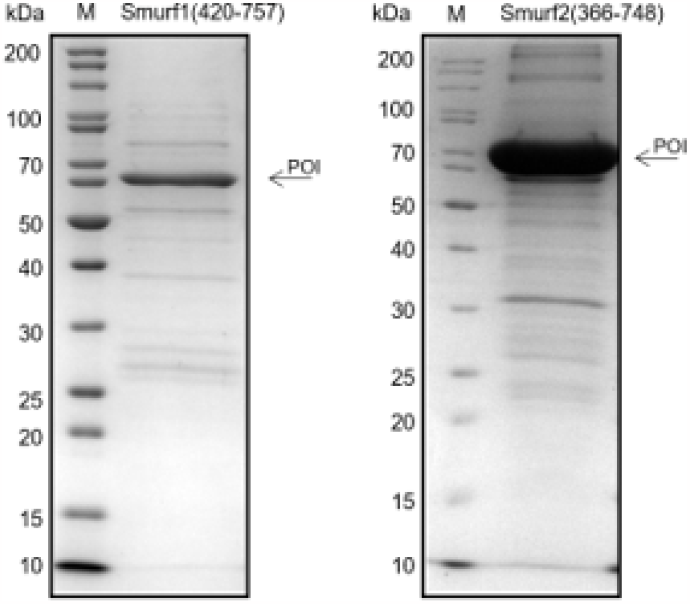
Truncated versions of GST-Smurfl(FL) and GST-Smurf2(FL) show higher purification yields compared to full-length proteins. Soluble fractions of lysates were subjected to affinity-purification and normalized amounts of eluates were analyzed by SDS-PAGE.

### Small scale-purification of isolated HECT domains

Bacterial pellets obtained from the most promising expression conditions (considering the amount of Protein of Interest (POI) in the soluble lysate fraction) underwent small-scale affinity purifications, following a similar procedure as the tested full-length constructs.

Notably, the expression of HECT domains alone, devoid of the N-terminal extensions, yielded significantly higher quantities of purified protein. This enhancement was evident for both Smurf1 (where the amount of protein in the elution increased from undetectable levels to a substantial amount) and Smurf2 (where very low levels improved to a high concentration of recombinant protein in the eluate).

It is worth emphasizing that the focus of this expression optimization strategy presented here was on identifying conditions where the largest amount of POI is present in the soluble fraction. This choice is motivated by the fact that purification from inclusion bodies involves denaturation and refolding steps, which often necessitate extensive optimization and condition testing. Moreover, this method poses a higher risk of obtaining a protein that may not be physiologically or biophysically relevant, potentially being fully or partially unfolded and/or prone to aggregation.

## Conclusions

A comparative analysis of expression conditions for three HECT E3 ligases revealed a consistent trend favoring lower expression temperatures. This preference is attributed to the positive impact of lower temperatures on solubility challenges associated with the Protein of Interest (POI). Lower temperatures serve as a tactical approach to enhance the solubility of the expressed HECT ligases, by preventing soluble and insoluble aggregation or protein misfolding. This in turn allows for efficient and soluble expression of HECT E3 ligases.

Based on the above-presented results, isolated HECT domains generally exhibit a higher propensity for successful expression in comparison to full-length HECT E3 ligases, although the difference in expression levels and yields of protein available for purification depends on the specific characteristics of the individual E3 ligase. In many instances, isolating and expressing the HECT domain alone can mitigate challenges associated with the full-length protein, leading to improved solubility and higher yields. However, it is important to note that the ease of expression can vary among different HECT ligases, and careful consideration of each ligase’s unique attributes is crucial for optimizing expression strategies tailored to their specific structural and functional features. Since studies of full-length E3 ligases is often necessary in biomedical research, future studies may focus on expression of HECT-type E3 ligases engaged in covalent complexes with ubiquitin as exemplified in literature. This could act as generalizable platform for production of these important proteins.

## Acknowledgements

We would like to thank Krzysztof Gumowski for the involvement in the project realization. This work was supported by the National Centre for Research and Development (NCBiR, Poland) project grant no. POIR 04.01.02-00-0147/16.

